# Response representation shapes AlphaGenome enhancer-gene prediction in human astrocytes

**DOI:** 10.64898/2026.08.14.744988

**Authors:** Zhongwei Huang, Ruohan Huang, Jinquan Han

## Abstract

**Background:** Sequence-to-function models predict regulatory perturbation responses, but their added value beyond established enhancer-gene predictors and across prediction tasks remains unclear. We evaluated whole-enhancer deletion and sparse variant responses in human astrocytes.

**Results:** We analyzed 2,307 CRISPR-interference-tested relations, including 133 expression-decreasing hits. Adding deletion responses increased fully nested EGrf average precision from 0.5475 to 0.6137. The paired increment was 0.0662 (enhancer-cluster 95% interval 0.0144-0.1181). In a separate exploratory analysis, sparse Atlas responses were combined with seven-input logistic and random-forest comparators by equal-weight rank fusion. Atlas magnitude fusion exceeded absolute-deletion fusion by 0.0519 and 0.0637 in average precision. Enhancer-cluster 95% intervals were 0.0098-0.0957 and 0.0220-0.1094, respectively. Standalone Atlas magnitude reached average precision 0.5570, but its advantage over absolute deletion remained uncertain. Signed Atlas fusion performed worse than either comparator alone. Incremental value for within-enhancer target choice and in the external WTC11 MHC evaluation remained uncertain. The reported performance intervals use one-way clustering of fixed predictions; Atlas analyses were exploratory within the same cohort.

**Conclusions:** In AstroREG, whole-enhancer deletion responses increased observed retrieval performance beyond a fully nested chromatin-based predictor. Sparse variant responses were more informative as magnitudes than as signed averages. Improved within-enhancer target choice and external transfer were not established.

## Background

Assigning distal regulatory elements to their target genes requires more than identifying active enhancers. Activity-by-contact (ABC) combines enhancer activity with contact information [1], while CRISPR interference (CRISPRi) screens measure which genes respond to repression of a tested element [2-4]. These experimental maps support supervised relation predictors and provide a demanding test for information derived from DNA sequence.

Sequence-to-function networks predict regulatory readouts from DNA, progressing from chromosome-scale convolutional models [5] to long-context models such as Enformer [6], Borzoi [7] and AlphaGenome [8]. However, accurate promoter-proximal predictions can coexist with weak distal-enhancer sensitivity [9,10]. A perturbation score might therefore discriminate CRISPRi hits yet add little beyond expression, distance or chromatin. Testing incremental value requires a comparator trained and evaluated without sharing target genes or enhancers across training and test sets [11].

The response representation creates a second uncertainty. Whole-enhancer deletion produces a directional response to one large edit; AlphaGenome Atlas supplies precomputed responses to individual substitutions [12]. Averaging signed effects asks whether local variants agree in direction, whereas averaging their magnitudes measures sensitivity regardless of direction. We therefore compared the two representations using the same absolute-value transformation and combination rule.

We examined these questions using AstroREG, a resource generated by CRISPRi screening of 979 candidate distal enhancers in primary human astrocytes, with an accompanying random-forest relation predictor, EGrf [4]. We first tested deletion responses against measured context and a fully nested EGrf reconstruction. We then compared sparse Atlas and deletion representations through matched rank fusion with separate pre-assay-feature comparators. Finally, we distinguished retrieval across the cohort from ranking candidate genes within an enhancer, and assessed supporting and external evidence without assuming transfer between these tasks.

## Results

### Deletion responses discriminate expression-decreasing relations

The evaluation comprised 2,307 relations across 745 enhancers and 1,054 genes, including 133 expression-decreasing hits (Figure 1a). The 2,174 comparison relations were well-powered tests without a source-defined expression-decreasing hit, including six expression-increasing hits. Each enhancer was deleted in silico once to score all eligible target genes. Deletion strength measured the predicted decrease in target-gene RNA after that edit.

**Figure 1.**
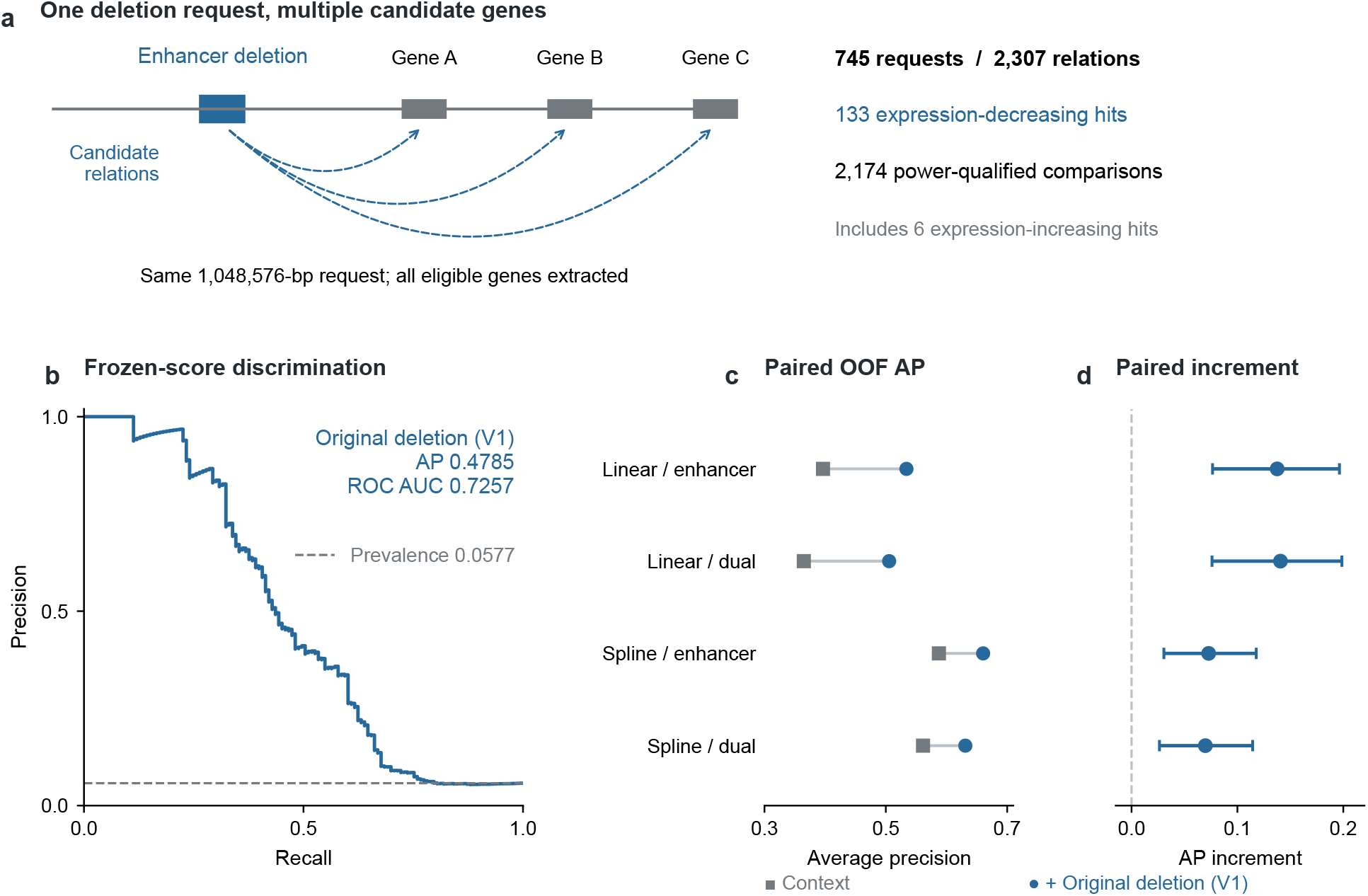
AstroREG request design, discrimination and paired context comparison. **a**, One enhancer deletion supplies scores for multiple candidate genes from the same 1,048,576-bp request; dashed arrows represent candidates, not established causal links. The cohort contains 745 requests and 2,307 relations: 133 expression-decreasing hits and 2,174 power-qualified comparisons, including six expression-increasing hits. **b**, Full empirical precision-recall curve for frozen V1 deletion strength; the dashed line shows prevalence. **c,d**, Aligned views of the same four context/holdout specifications: paired out-of-fold AP (gray squares, context; blue circles, context plus AlphaGenome) and paired AP increments with 95% enhancer-cluster intervals from 20,000 resamples. “Dual” denotes crossed gene-enhancer holdout; “spline” denotes nonlinear measured-expression adjustment.

In the prespecified primary deletion analysis, average precision (AP) was 0.4785 against prevalence 0.0577 (enhancer-cluster 95% interval 0.3915-0.5650), with ROC AUC 0.7257 (Figure 1b). A post-hoc sensitivity excluding relations closer than 10 kb retained discrimination (AP 0.420; prevalence 0.045; Supplementary Table 3a). Distance and ABC standalone scores provided additional reference points, with APs of 0.3075 and 0.4013, respectively (Supplementary Table 17).

### Deletion responses improve retrieval in AstroREG beyond context and nested EGrf

We first compared models containing distance, ABC score, enhancer length, measured expression and assayed-cell count. These variables include expression in guide-bearing cells and describe experimental context. Adding deletion strength increased enhancer-held-out AP from 0.3965 to 0.5340 (paired increment 0.1375, 95% interval 0.0765-0.1964) and reduced log loss from 0.1688 to 0.1497. AP and log-loss gains persisted with nonlinear expression adjustment and crossed gene/enhancer holdout (Figure 1c,d; Supplementary Tables 1 and 10). Matching without AlphaGenome outputs also retained a positive score difference, although distance, ABC and expression remained imperfectly balanced (Supplementary Table 6).

EGrf offered a stronger comparator using seven chromatin and relation features from the same resource. We reconstructed it with tuning, forest fitting and calibration confined to each outer training partition. Across 25 crossed gene/enhancer test cells, calibrated EGrf achieved AP 0.5475; adding deletion strength increased AP to 0.6137 (Figure 2a). The paired increment was 0.06624 (enhancer-cluster 95% interval 0.01445-0.11808). Log loss improved by 0.01039, whereas the ROC-AUC increment remained uncertain (Figure 2b; Supplementary Table 7). At a retrospective budget of 100 relations, precision increased from 0.56 to 0.69, corresponding to 13 additional known hits in this completed cohort. Precision at 25 remained 0.92 (Figure 2c; Supplementary Table 4).

**Figure 2.**
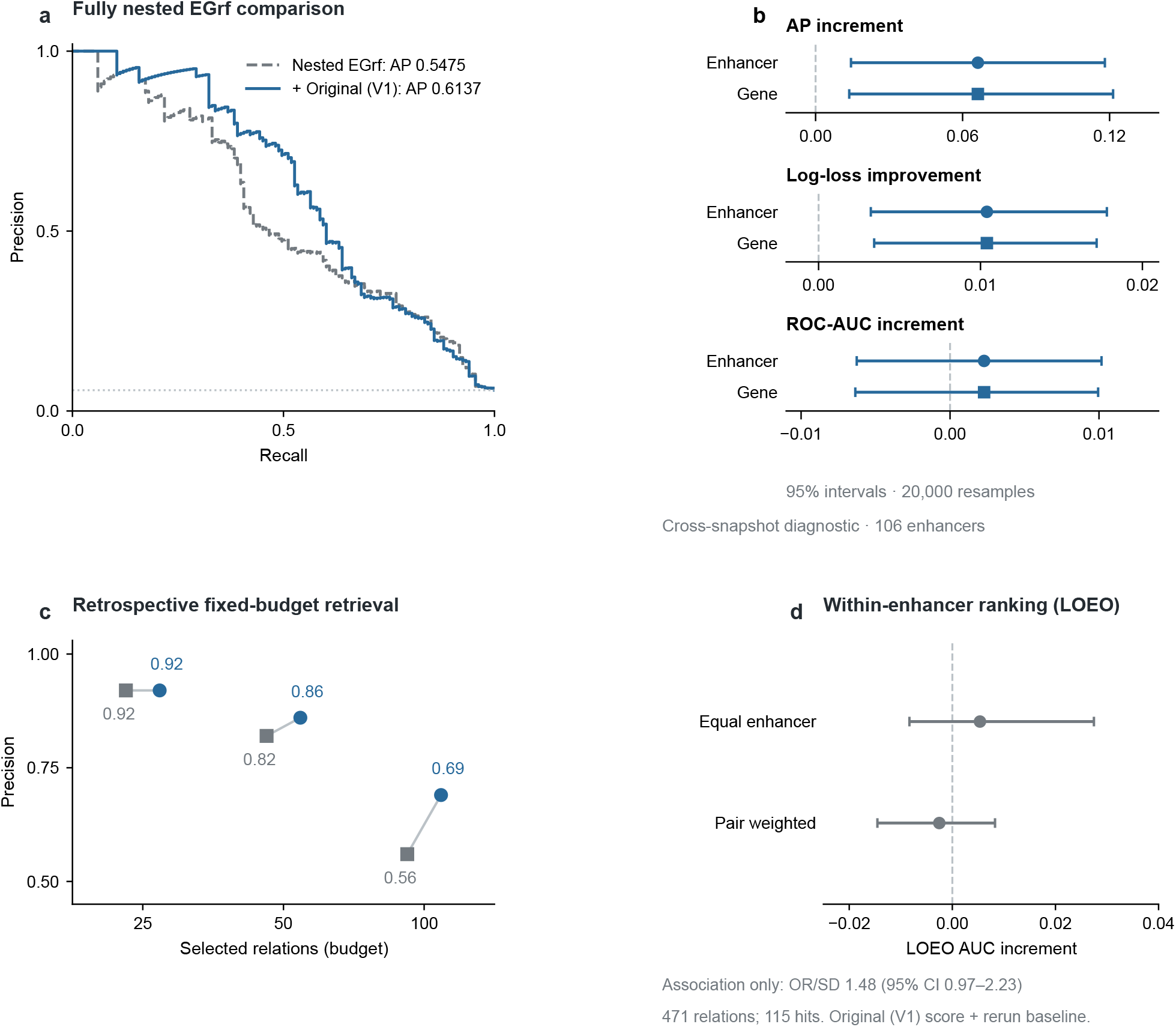
Fully nested EGrf comparisons and within-enhancer ranking. **a**, Full empirical precision-recall curves from 2,307 outer out-of-fold relations (133 hits): calibrated nested EGrf (gray dashed) and its V1 AlphaGenome stack (blue solid). The dotted line shows prevalence. **b**, Paired AP and ROC-AUC increments and log-loss improvement on separate natural-unit axes. Circles and squares show enhancer-cluster and gene-cluster 95% intervals, respectively, each using 20,000 resamples. **c**, Retrospective precision for the top 25, 50 and 100 relations. Gray squares and blue circles compare the same models within each budget. Precision at 25 is unchanged. **d**, Leave-one-enhancer-out (LOEO) adjusted AUC increments within 106 mixed-outcome enhancers (471 relations, 115 hits), with two-sided 95% intervals from 20,000 enhancer resamples. This cross-snapshot diagnostic combines V1 deletion scores with rerun baseline activity. The secondary OR annotation describes association, not predictive gain.

The deletion-associated AP increase persisted after adjustment for the model’s predicted baseline expression of the target gene in the reported one-way analyses. Adding matched rerun deletion to nested EGrf plus predicted baseline increased AP from 0.5411 to 0.6053 (increment 0.06422, enhancer-cluster interval 0.01181-0.11623). Context-model gains also persisted after restricting requests to contained exon unions or interior target transcription start sites (Figure 3; Supplementary Tables 2, 3b and 7). Length-preserving dinucleotide shuffling retained discrimination; under signed scoring, middle-third deletion reduced it relative to full deletion (Supplementary Table 18). These controls address baseline activity, request geometry and perturbation encoding without isolating a single causal explanation.

**Figure 3.**
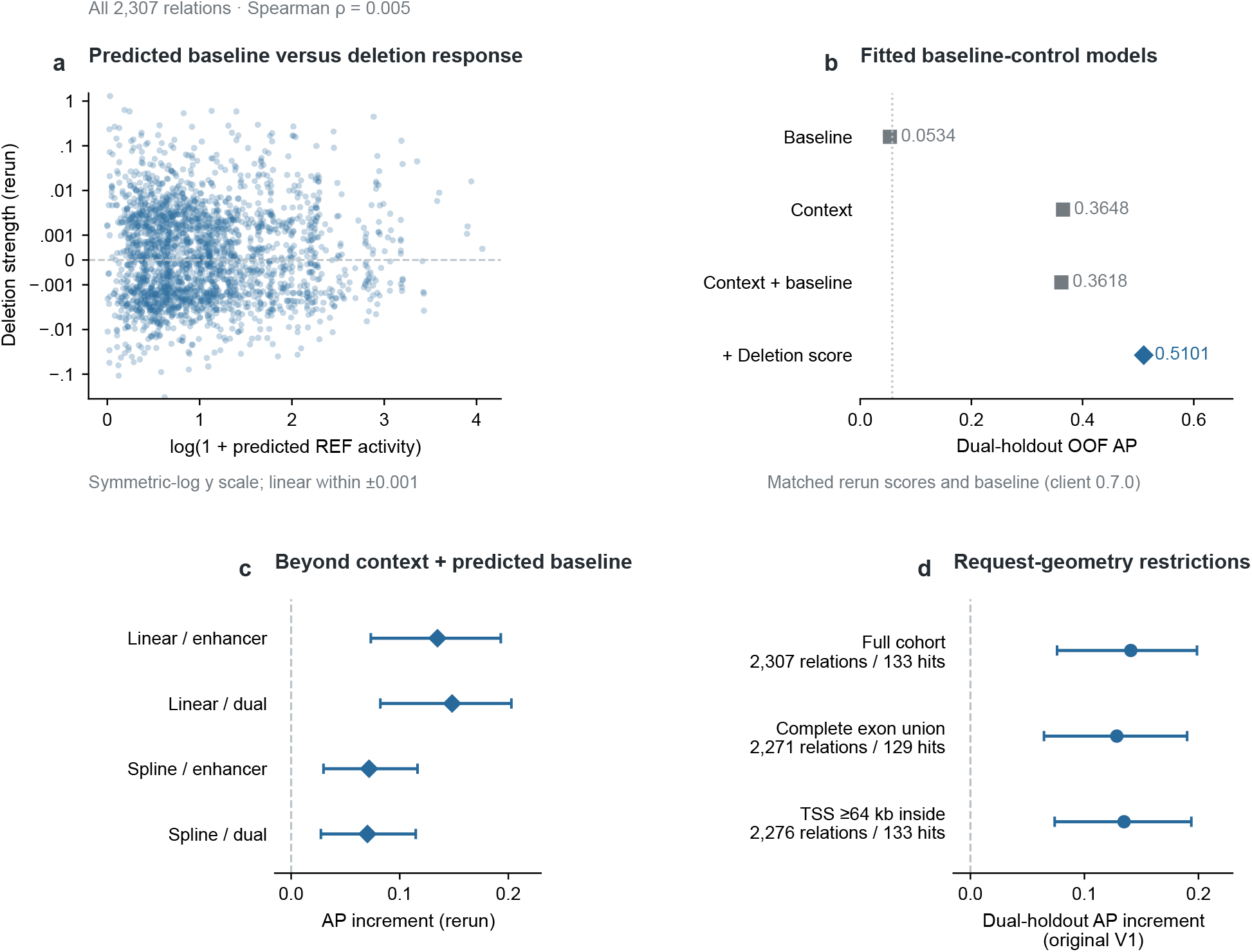
Version-matched baseline controls and request-geometry sensitivities. **a**, All 2,307 rerun deletion scores versus log-transformed reference activity, with full ranges preserved. The symmetric-log vertical axis is linear within ±0.001. Near-zero Spearman correlation does not establish independence. Rerun scores used client 0.7.0. **b**, Linear dual-holdout out-of-fold AP for fitted baseline, context, context-plus-baseline and context-plus-baseline-plus-deletion models; the final model is a blue diamond and comparators are gray squares. The dotted line is prevalence. Fitted baseline-only AP 0.0534 differs from raw baseline-score AP 0.0694 (Supplementary Table 2). **c**, Increment from adding matched rerun deletion scores beyond context and rerun baseline. **d**, V1 increments beyond linear dual-holdout context in the full cohort and two request-geometry subsets; relations/hits identify each population. Bars in c,d are 95% enhancer-cluster intervals from 20,000 resamples. Point APs and scatter associations have no plotted intervals.

### Sparse variant magnitude under equal-weight rank fusion

Atlas provided RNA effects for all three alternate bases at three enhancer positions, yielding nine substitutions per relation and 20,763 variant-gene responses [12]. We summarized the same effects by mean absolute magnitude or negative mean signed effect. Standalone Atlas magnitude achieved AP 0.5570 and ROC AUC 0.8712, compared with 0.5018 and 0.8581 for absolute rerun deletion. Their paired AP difference was uncertain (0.0552, enhancer-cluster 95% interval -0.0098 to 0.1225). Signed Atlas had AP 0.2666 and ROC AUC 0.5232 (Supplementary Table 17).

We combined response ranks with separately fitted seven-input logistic and random-forest comparators, using fixed equal weights and crossed gene/enhancer holdout (exploratory; Methods). The inputs were available before perturbation and excluded guide-bearing expression and assayed-cell count. Because these comparators differ from the ranger EGrf reconstruction, comparisons of response partners were made within this pipeline.

Atlas magnitude fusion achieved AP 0.6155 for logistic regression and 0.6357 for random forest, compared with comparator-only ranks of 0.5110 and 0.5157. Absolute-deletion fusion achieved 0.5636 and 0.5720 under the same rank map and weights (Figure 4a). Direct Atlas-minus-deletion increments were 0.0519 (enhancer-cluster 95% interval 0.0098-0.0957) and 0.0637 (0.0220-0.1094; Figure 4b). The point estimates favor Atlas as the fusion partner under this rule. The Atlas-deletion comparisons replace the response partner; they do not test adding Atlas to a model already containing deletion. Perturbation type and request geometry also differ between response scores.

**Figure 4.**
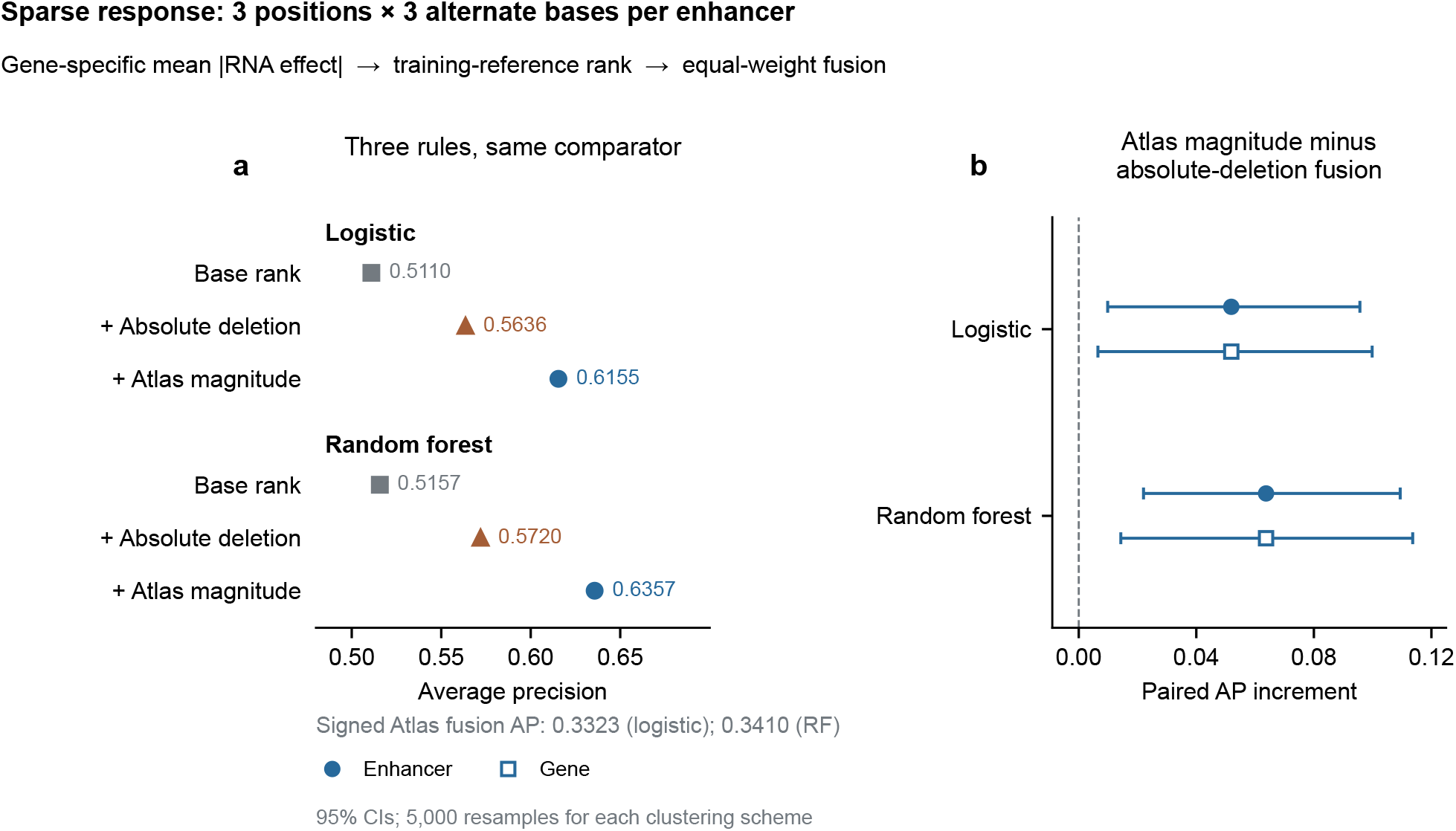
Atlas magnitude and absolute deletion under matched rank fusion. **a**, Average precision for the base comparator rank, absolute-deletion fusion and Atlas-magnitude fusion within each seven-input comparator. Values are point estimates for the same 2,307 relations (133 hits); rank scores are not calibrated probabilities. Atlas summarizes three positions and three alternate bases per position. The annotation retains the adverse signed-decrease fusion results. **b**, Direct paired Atlas-minus-absolute-deletion fusion increments. Circles and squares show 95% percentile intervals from 5,000 enhancer- or gene-cluster resamples of the same fixed predictions. The exploratory, outcome-informed analysis is detailed in Supplementary Tables 11-12, including ROC-AUC contrasts.

Signed Atlas fusion reached APs of 0.3323 and 0.3410, below both comparators alone (Figure 4; Supplementary Table 11). Thus, magnitude was more useful than the signed summary tested here. ROC-AUC differences between Atlas-magnitude and absolute-deletion fusion remained uncertain (Supplementary Table 12). An enhancer-level AVI summary did not show a clear AP increment when added to either comparator (Supplementary Table 19).

### Within-enhancer rankings and increments over proximity

Retrieving responsive relations across a cohort differs from ranking candidate genes for a particular enhancer. Among 106 mixed-outcome enhancers (471 relations; 115 hits), adding deletion to an adjusted leave-one-enhancer-out model gave an equal-enhancer AUC increment of 0.00536 (95% interval -0.00832 to 0.02746; Figure 2d). This cross-snapshot diagnostic and the snapshot-matched association checks are distinguished in Methods and Supplementary Tables 5 and 13.

Atlas magnitude and absolute rerun deletion ranked candidate genes with equal-enhancer AUCs of 0.8965 and 0.8861, compared with 0.8382 for proximity. The Atlas-deletion difference was 0.0105 (95% interval -0.0451 to 0.0648), and the Atlas-proximity difference was 0.0584 (-0.0094 to 0.1250). The secondary pair-weighted Atlas-proximity difference was positive (0.1215, 0.0230-0.2086), showing dependence on how enhancers contribute to the summary (Supplementary Table 14). These scores rank candidate genes, but an increment over proximity was not established for the primary equal-enhancer endpoint.

### External evaluation against rE2G

The WTC11 iPSC MHC CRISPRi evaluation [13] contained 317 cis-regulatory elements (CREs) and 5,557 relations. Standalone Atlas magnitude had mean within-CRE Spearman correlation 0.1162, compared with -0.0067 for signed Atlas and 0.1142 for rE2G [3,14]. Signed fusion reduced correlation relative to rE2G by 0.0480 (95% interval -0.0761 to -0.0200); magnitude fusion increased it by 0.0155 (-0.0074 to 0.0372). Neither fusion established a positive increment (Supplementary Table 15). This one-donor, one-locus test used within-CRE ranks rather than transferring the AstroREG classifier.

The supporting K562 analysis was tail-sensitive and overlapped released benchmarks, so it provides supporting rather than independent evidence (Supplementary Figure 1; Supplementary Tables 8-9).

## Discussion

The value of a sequence-model response depends on its representation, the information it complements and the prediction task. In AstroREG, adding deletion strength increased the observed AP of context models and nested EGrf. Atlas magnitude yielded higher AP than absolute deletion under a separate equal-weight fusion rule. The contrast with signed Atlas and the within-enhancer results shows why response representation and evaluation task should be considered together.

AlphaGenome already supported enhancer-gene benchmarks [8,15], and Atlas introduced genomic-scale variant-response resources [12]. Our contribution is to test added value against nested in-domain predictors and to separate response representation from prediction task. The Atlas comparison holds the absolute-value transformation, comparator and fusion weights constant. Atlas had the higher AP point estimate under the matched equal-weight fusion rule. Added value beyond a combined deletion-plus-comparator model remains untested. Sparse substitutions and whole-enhancer deletion also differ in perturbation type and request geometry.

The contrast between magnitude and signed averaging changes the interpretation of sparse variant responses. Magnitude preserves sensitivity when effects of opposite sign cancel, which may explain its better performance here. Earlier evaluations of personal gene-expression prediction also found difficulties recovering effect direction [16,17], although those studies tested different models and tasks. Our results favor the magnitude summary tested here; they do not establish that direction is generally uninformative or identify a biological mechanism.

Within-enhancer ranking and cohort retrieval answer different prioritization questions. Atlas magnitude ranked candidate genes well, but its equal-enhancer increment over proximity remained uncertain; pair-weighted results were more favorable. Consequently, good pooled retrieval does not by itself establish additional value for selecting the target of an individual enhancer. Practical evaluation should compare signed and magnitude summaries, test added value against a strong comparator, and report within-enhancer performance separately. Recording score versions is also necessary: original and rerun deletion responses had Spearman correlation 0.831 (Supplementary Methods, AlphaGenome client-0.7.0 rerun).

### Limitations of the study

AstroREG’s expression and power filters define the tested population, and its public release preceded AlphaGenome. Pre-assay comparators still require chromatin assays, and nested fitting does not undo their historical feature selection. The Atlas analyses followed inspection of AstroREG outcomes; Methods distinguishes the sampling plan from later fusion and magnitude-control choices. The reported one-way intervals do not jointly account for gene and enhancer dependence, so they do not establish robustness to that crossed structure. Performance intervals based on fixed predictions omit model-refitting and analysis-selection uncertainty. The MHC and K562 results do not establish broad transfer. An external evaluation fixing the representation and comparator before examining outcomes would test whether the retrieval rule generalizes beyond this cohort.

## Conclusions

In AstroREG, whole-enhancer deletion responses increased AP and reduced log loss beyond a fully nested chromatin-based predictor. Sparse variant responses were more informative as magnitudes than as signed averages under the tested fusion rule. These findings concern retrieval within the evaluated cohort and the reported uncertainty analyses; improved within-enhancer target choice and external transfer were not established.

## Methods

### Study design and analysis status

All analyses were computational re-analyses of public experimental data. In the original deletion study, the AstroREG cohort, AlphaGenome output, deletion encoding, primary dependence unit, discrimination metrics, context variables, grouped cross-validation and matching without AlphaGenome outputs were frozen before AstroREG AlphaGenome outputs were inspected. The baseline, geometry, fully nested EGrf and adjusted same-enhancer modules were subsequent analyses designed to test specified alternative explanations. They were not used to redefine the cohort. The K562 analysis was frozen earlier and is secondary here. The Atlas analyses were exploratory and began after AstroREG outcomes were known. Nine-substitution sampling and mean-absolute and signed summaries were specified before cohort-wide retrieval. Fusion rules were fixed after standalone Atlas results were seen; the absolute-deletion control followed the first fusion results. These local plans document analysis order, not external preregistration. Supplementary Information records the analysis stages and source configurations.

### AstroREG cohort and outcomes

AstroREG source tables accompanied Green et al. [4] and use hg38 coordinates. We joined source records by the authors’ relation identifier. An expression-decreasing hit required the source permissive negative-direction hit indicator. A comparison relation required absence of that indicator and at least 80% power to detect a 15% effect according to the author-provided power table. The supplied cohort already contained unique relation identifiers and unique enhancer-gene pairs; no additional duplicate removal was required. The final cohort contained 133 expression-decreasing hits and 2,174 power-qualified comparison relations across 745 enhancers and 1,054 Ensembl genes. Of the comparison relations, 2,168 were not source-defined significant hits and six were significant expression-increasing hits (three core and three permissive-only). All 2,174 met the source power threshold. The endpoint is therefore the source-defined expression-decreasing class versus its power-qualified complement; the comparison class is not a set of biologically inactive relations.

The third-party AstroREG table is not redistributed because its source repository did not declare a repository-wide license at audit. A pinned retrieval script downloads the exact source file from the source revision specified in the accompanying version manifest and joins it to the derived score table. Relation identifiers, row counts and outcome counts are validated during reconstruction.

### AlphaGenome scoring

The scoring configuration used AlphaGenome GeneMaskLFCScorer(OutputType.RNA_SEQ) and a primary-astrocyte total RNA-seq output (CL:0000127). Each tested enhancer interval was represented as a VCF-compatible deletion anchored by the immediately preceding reference base. A 1,048,576-bp interval was selected to contain the enhancer and all eligible target-gene transcription start sites. One API request was issued for each enhancer, and every eligible Ensembl target gene was extracted from the same response. Deletion strength was the negative ALT-versus-REF gene-mask RNA-seq log-fold change. No missing score was converted to zero.

The principal analysis preserves the original (V1) AlphaGenome outputs. A separate complete rerun used client version 0.7.0 and the then-current server default to obtain internally version-matched deletion and predicted-reference quantities. Rerun outputs were not substituted for original V1 values. The public API did not expose a server snapshot identifier sufficient to reproduce the historical V1 default.

### Primary discrimination and clustered uncertainty

AP was the primary discrimination measure because positives comprised 5.77% of relations; ROC AUC was secondary. The raw-score analysis used 20,000 percentile bootstrap resamples of enhancer blocks. Gene-cluster resampling assessed sensitivity to target-gene dependence; it does not itself establish generalization. Mean score separation compared expression-decreasing hits with power-qualified comparison relations. Direction was descriptive because the positive outcome definition selected negative expression effects.

### Context models and dual holdout

Context variables were log1p absolute distance, log10(ABC + 0.000001), log1p enhancer length, log1p measured expression and log1p assayed-cell count. Logistic regressions used training-fold standardization and effectively unpenalized fitting (C=1,000,000). The linear specification used measured expression as one term. The flexible specification transformed expression with a cubic spline using four quantile knots fitted only in training data.

Enhancer-held-out predictions used five GroupKFold partitions. Dual holdout crossed five gene folds with five enhancer folds; for each test cell, training excluded every row belonging to either held-out group. The principal deterministic layout used seed 20260828 where random assignment was required. Three alternate stratified-group layouts used seeds 20260829-20260831. Paired AP, ROC-AUC and log-loss differences used 20,000 enhancer-cluster resamples of the completed out-of-fold predictions.

The source measured-expression covariate is the mean normalized RNA expression across guide-bearing cells, not a separately assayed control-only unperturbed baseline; assayed-cell count is the enhancer guide-positive cell count. These are experimental-context covariates, not inputs for sequence-only deployment.

### Predicted-baseline activity control

Rerun predicted reference activity was reconstructed from the matched scorer outputs and log-transformed. Its relation-level association with rerun deletion strength was summarized by Spearman and Pearson correlations. We compared baseline alone, context, context plus baseline, and context plus baseline plus rerun deletion strength using the same grouped folds and metrics. The historical baseline is unavailable. The baseline-control models and the corresponding nested EGrf stack use rerun deletion and rerun baseline quantities together; the secondary adjusted same-enhancer diagnostic is distinguished below.

### Gene-mask and request-window geometry

The official AlphaGenome GENCODE v46 hg38 annotation defined gene bodies and the union of annotated exons. Coordinates were treated as 0-based, half-open intervals. We calculated the fraction of each target exon union within the original request, complete exon-union and gene-body containment, and distance from the target transcription start site to the nearest request boundary. Two bounded sensitivities were evaluated: complete exon-union containment and at least 64 kb between the transcription start site and either request boundary. The analysis tests geometric truncation; it is not an exact reimplementation of undisclosed server-side transcript selection.

### Fully nested EGrf reconstruction

We mapped the seven released EGrf features exactly: H3K4me3, H3K27ac, nearest-gene status, absolute distance, ATAC-seq pileup, number of transcription start sites between enhancer and gene, and gene-stability index. The author probability was evaluated descriptively because its hyperparameters were tuned before author cross-fitting.

The nested reconstruction used ranger probability forests. The outer split crossed the same five gene folds and five enhancer folds used for dual holdout. Within each outer training partition, a 3-by-3 grouped split selected among 12 combinations: mtry 2, 3 or 5; Gini or extremely randomized splitting; and minimum node size 1 or 10. Tuning forests used 200 trees; the selected final forest used 1,000 trees. Hyperparameters were selected by inner out-of-fold AP, with log loss and fixed parameter order as tie-breakers. Logistic calibration of EGrf alone, EGrf plus V1 AlphaGenome, EGrf plus rerun baseline, and EGrf plus rerun baseline plus rerun AlphaGenome was trained only on inner out-of-fold predictions from the corresponding outer training partition. All reported predictions were outer out-of-fold. Paired differences used 20,000 enhancer- and gene-cluster resamples.

### Adjusted same-enhancer association and predictive ranking

Mixed-outcome enhancers contained at least one expression-decreasing hit and one power-qualified comparison relation. The fixed population comprised 106 enhancers, 471 relations and 115 hits. Conditional logistic regression stratified by enhancer and used standardized V1 AlphaGenome deletion strength, log distance, log ABC, log measured expression, log rerun predicted baseline and GENCODE exon-union coverage. The baseline is an auxiliary covariate from a later snapshot, so this is a cross-snapshot association and ranking diagnostic, not evidence of version-matched baseline independence. We report the AlphaGenome odds ratio per standard deviation, an enhancer-cluster sandwich interval, robust Wald P value and a one-degree-of-freedom likelihood-ratio test against the same model without AlphaGenome.

A snapshot-matched learned leave-one-enhancer-out comparison was not performed. For predictive ranking, models were refitted while holding out one entire enhancer at a time; scaling parameters were learned on training enhancers only. Within each enhancer, AUC was the proportion of hit-comparison pairs correctly ordered, with half credit for ties. We report equal-enhancer and comparison-pair-weighted summaries. Their paired increments used 20,000 enhancer bootstrap resamples. Candidate multiplicity was fixed as 2, 3-4 or at least 5 tested genes.

### Matching without AlphaGenome outputs

The frozen match paired each expression-decreasing hit without replacement to a comparison relation by minimizing total pairwise Euclidean distance in standardized log distance, log ABC, log enhancer length, log measured expression and log assayed-cell count, without using AlphaGenome output. We calculated pre- and post-match standardized mean differences and common support. A single sensitivity retained pairs below the 95th percentile of the original matching distance. Mean paired differences used 20,000 enhancer-cluster resamples.

### Fixed-budget ranking

Precision was calculated among the top 25, 50 and 100 out-of-fold relations. Recall was calculated in the top 1%, 5% and 10% of the completed cohort. Budgets were fixed before calculation and describe ranking within the completed cohort.

### Supporting K562 analysis

K562 records were derived from a DNALONGBENCH-formatted Gasperini CRISPRi table [2,10]. The frozen cohort paired 161 functional regions with comparison regions within 121 target genes. Gene-equal mean, trimmed, winsorized and median contrasts used 20,000 gene-level resamples. Influence analyses removed the largest positive or largest absolute contrasts only as descriptive stress tests. Exact gene-and-interval joins quantified overlap with released AlphaGenome K562 benchmark rows [15] and ENCODE-rE2G source tables [3]. These audits determined the supporting status of K562; they did not alter the AstroREG cohort or result.

### Atlas sampling and score identity

The expansion accessed Atlas using client 0.9.0 on 11 September 2026 [12,18]. For a zero-based enhancer interval [start,end), positions were floor(start + q^*^(end-start)), for q=0.25, 0.50 and 0.75. All three alternate substitutions were queried at each position. We used raw signed RNA_SEQ effects from CL:0000127 total RNA-seq, not active-allele or quantile scores. Stable Ensembl identifiers were joined after resolving aliases against GENCODE v46. Mean absolute and negative mean signed effects used the same nine substitutions. All 26 initial identifier omissions were systematically recovered; no final missing response was imputed.

Atlas uses 128-bp block-centered, 1,048,576-bp windows, whereas whole-deletion requests were designed around an enhancer and its candidate genes. Window and mask differences remain part of the comparison. The client-0.9.0 whole-deletion evaluation reproduced the cached client-0.7.0 rerun scores in this cohort; the hosted service did not expose a weight revision. V1, the client-0.7.0 rerun and the 11 September Atlas access remain distinct identities. Figures distinguish original (V1) scores from rerun scores.

### Pre-assay comparators and fixed fusion

The seven inputs were H3K4me3, H3K27ac, ATAC pileup, nearest-gene status, absolute enhancer-TSS distance, intervening TSS count and an external gene-stability index. They were retained from the pinned AstroREG table. Stability zeros, used upstream as a missing-lookup fallback, were treated conservatively as zero-or-unavailable in 143 relations and handled by training-median imputation plus an indicator. Literal missing values followed the same training-only procedure; ChIP zero means were retained. Guide-bearing expression, cell counts, ABC and predicted reference activity were excluded.

Outer folds crossed the existing five gene and five enhancer partitions, excluding both held-out groups from training. Inner tuning crossed three gene and three enhancer groups inside each outer training set. Logistic regression used log1p on nonnegative chromatin, distance and TSS-count inputs, raw nearest-gene and stability inputs, training-only scaling, and C in {0.1,1,10}. Random forests used untransformed inputs, max_features=3, min_samples_leaf in {1,5,20}, 200 inner trees and 500 final trees. Selection maximized inner out-of-fold AP, with log loss and grid order breaking ties. These implementations differ from ranger EGrf.

For reference distribution T, the mid-CDF was r_T(s)=(count(t<s)+0.5^*^count(t=s))/|T|. Comparator test predictions used the selected model’s inner out-of-fold predictions as T; response scores used the corresponding outer-training response values. Fusion was the arithmetic mean of the two ranks, compared with an identically mapped comparator-only rank. No RNA output entered predictive fitting, hyperparameter selection or calibration. Equal weighting provided a common combination rule without estimating response-specific weights. The analysis order is reported under Study design and analysis status.

Atlas paired contrasts used 5,000 enhancer-cluster and 5,000 gene-cluster percentile resamples of fixed out-of-fold predictions. The two one-way procedures address alternative dependence structures; neither jointly resamples genes and enhancers. Historical deletion analyses retain their 20,000-resample procedures. Performance intervals computed from fixed predictions omit model-refitting and analysis-selection uncertainty.

### External continuous endpoint and secondary analyses

The WTC11 common-support set contained 5,591 relations, 328 CREs and 53 genes after previously documented geometry, finite rE2G/guide-effect and old K562-overlap restrictions. Exactly 5,590 relations had nine finite scores. One relation had six because one window excluded its target TSS and was left missing. Requiring at least four candidates yielded the 317-CRE primary endpoint. RNA track EFO:0009747 matched WTC11 identity, without establishing personalized haplotypes or identical culture conditions.

The outcome was negative median published guide log2 fold change. Fusion averaged within-CRE percentile ranks with average ties. Mean within-CRE Spearman used 10,000 CRE-summary resamples. The separate equal-gene endpoint used a correspondingly transformed rE2G-only comparator because within-CRE normalization can change across-CRE ordering. This is not literal transfer of the AstroREG model. Complete-score and full-span sensitivities, secondary FRACTEL outcomes, scalar AVI controls and conditional association details are retained in Supplementary Tables 13-16 and the accompanying source tables.

### Software and reporting

Core analyses used Python 3.11 with NumPy, pandas, SciPy and scikit-learn. Nested EGrf used R and ranger; conditional regression used R survival. Figures were generated with Matplotlib and exported as editable SVG, PDF and 600-dpi LZW-compressed TIFF. Machine-readable predictions, metric tables, resampling counts, selected hyperparameters and source versions are retained in the publicly archived reproducibility package (version 1.0.0; https://doi.org/10.5281/zenodo.23351607). Statistical inference used clustered resampling or model-based procedures; nominal P values in the adjusted same-enhancer module are not promoted to confirmatory evidence.

### Use of generative AI

Generative-AI tools, including OpenAI Codex, Claude and DeepSeek, assisted selected aspects of code development, literature triage, analysis planning and auditing, and manuscript drafting and editing.

## Abbreviations

ABC: activity-by-contact
AP: average precision
AUC: area under the curve
CRE: cis-regulatory element
CRISPRi: CRISPR interference
OOF: out-of-fold
TSS: transcription start site.

## Declarations

### Availability of data and materials

Code, frozen predictions, source tables and figure-generation scripts are available at https://github.com/booksiyideEdward/op26-alphagenome-astroreg and https://doi.org/10.5281/zenodo.23351607 (version 1.0.0). The Atlas extension is under atlas_extension/; versions and provenance are in docs/ATLAS_VERSION_AND_PROVENANCE.md. Offline checks verify fixed-score metrics and fusion identities. The reconstruction script retrieves the pinned AstroREG author table from its official source. Complete third-party source tables, model weights and credentials are excluded. AlphaGenome outputs and derivatives remain governed by applicable service/output terms, not the MIT code license. K562/AstroREG sources are identified above; MHC/rE2G resources are references 13-14.

### Ethics approval and consent to participate

This study performed secondary computational analyses of publicly released, de-identified experimental datasets and generated no new human-participant or animal data. Ethical approvals and participant-consent procedures for the source materials are described in the original studies.

### Funding

This research received no specific funding.

### Competing interests

The authors declare no competing interests.

### Authors’ contributions

**Zhongwei Huang:** Conceptualization, Methodology, Software, Data curation, Formal analysis, Investigation, Validation, Visualization, Writing - original draft, Writing - review & editing, Project administration. **Ruohan Huang:** Validation, Writing - review & editing. **Jinquan Han:** Software (code testing).

### Consent for publication

Not applicable. This manuscript contains no identifiable individual participant information.

## Acknowledgements

Not applicable.

## Notes

https://github.com/booksiyideEdward/op26-alphagenome-astroreg

https://doi.org/10.5281/zenodo.23351607

### Competing Interest Statement

The authors have declared no competing interest.

### Summary of Updates

This revision adds sparse AlphaGenome Atlas response analyses and a fully nested EGrf comparison. The supplementary material expands standalone, signed-versus-magnitude, matched rank-fusion and within-enhancer controls, and reports the external WTC11 MHC evaluation. The title, abstract, figures and discussion have been revised to distinguish cohort retrieval from within-enhancer target choice and to clarify the exploratory analysis and fixed-prediction, one-way clustering scope. The description of the original AstroREG screen has been corrected. Code, frozen predictions, source tables and reproducibility documentation are now publicly archived at https://doi.org/10.5281/zenodo.23351607.

