## Supplementary Methods, Tables 1-19 and Supplementary Figure 1 for "Response representation shapes AlphaGenome enhancer-gene prediction in human astrocytes"

#### Source reconstruction and relation integrity

The public reconstruction workflow (`scripts/prepare_journal_inputs.py --download-egrf`) downloads the AstroREG author table from commit `ec6e4e08f64b322935bc0dca2f5392ec1779b1a8` and validates its recorded checksum before joining it to `data/derived/astroreg_journal_relation_outputs.csv`. The join key is the unique author `Pair` identifier. The workflow requires 2,307 unique relations, 133 positive labels, 745 enhancers and 1,054 genes and rejects changes in the author gene identifier or outcome. The power-qualified comparison class includes 2,168 source-nonsignificant relations and six significant expression-increasing hits (all retained under the source-defined negative-direction endpoint). Author features and labels are kept in an ignored local analysis table because the source repository does not declare a repository-wide license; Project-derived scores and geometry fields remain public.

#### Cluster-resampled metric intervals

For relation-level discrimination, enhancer blocks were sampled with replacement, with the number of sampled blocks equal to the number of unique enhancers. Each sampled block carried all of its relations. Average precision, ROC AUC and log loss were recalculated using the resulting integer block weights. Paired increments used the same resample for the comparator and augmented prediction vectors. Gene-cluster intervals repeated this procedure with gene blocks. The original deletion/context/nested-EGrf, same-enhancer and supporting K562 bootstrap intervals use 20,000 resamples with fixed analysis-specific seeds. Atlas fixed-fusion intervals use 5,000 paired resamples; the external MHC endpoint uses 10,000 CRE-summary resamples. These module-specific counts are retained, not harmonized retrospectively. Primary raw-score intervals use seed 20260808; journal modules use base seed 20260828 with fixed module offsets recorded in their executable code; supporting K562 modules use seeds 20260808 and 20260809. Percentile limits are reported without a normal approximation.

The primary mean hit-minus-comparison deletion-strength difference was 0.08189 (enhancer-cluster 95% interval 0.05462-0.11402). Of the 133 expression-decreasing hits, 100 had a predicted decrease after deletion. Because the experimental positives were selected for decreased expression, this proportion is descriptive rather than an independent signed test.

#### Fold construction

The primary enhancer holdout used five deterministic group folds. The dual holdout independently assigned genes and enhancers to five folds and crossed them into 25 test cells. A relation entered a test cell only when both its gene and enhancer matched that cell. Its training data excluded every relation sharing either the held-out gene fold or the held-out enhancer fold. This produces smaller training partitions than ordinary fivefold cross-validation but prevents both forms of group overlap. Three alternate stratified-group assignments were evaluated once, with seeds 20260829, 20260830 and 20260831 (Supplementary Table 10).

#### AlphaGenome client-0.7.0 rerun

The rerun used AlphaGenome client 0.7.0, with the same ontology, RNA-seq output, relation identifiers, deletion coordinates and request construction as V1. All 745 requests and 2,307 relations completed. Rerun and V1 deletion scores were concordant (Spearman rho 0.8314; Pearson r 0.99945), but only 17 values agreed within 0.00001. Maximum and median absolute differences were 0.01164 and 0.000967, respectively. The source identifier “current” denotes this complete rerun; figures and tables label it “rerun”. The baseline-control models and corresponding nested EGrf stack pair predicted reference activity with deletion scores from that rerun. The adjusted same-enhancer model instead uses V1 deletion strength and rerun reference activity. Its cross-snapshot estimates are not a version-matched baseline-independence test, and the historical V1 reference activity remains unavailable.

#### Geometry annotation

The official AlphaGenome GENCODE v46 hg38 feather annotation was used. For each Ensembl gene, transcript exons were merged into a union before calculating covered bases and request containment. The analysis also recorded gene-body containment and the distance from the annotated transcription start site to the nearest request boundary. The exact annotation input is identified in the accompanying source-version manifest.

#### Fully nested EGrf tuning and stacking

Within each outer training partition, genes and enhancers were independently assigned to three inner folds and crossed into nine inner test cells. Twelve `ranger` candidates crossed `mtry` 2, 3 and 5; `gini` and `extratrees` split rules; and minimum node size 1 and 10. Each candidate generated complete inner out-of-fold probabilities using 200-tree forests. Selection maximized inner AP and then minimized log loss; deterministic parameter order resolved any remaining tie. A 1,000-tree forest fitted the complete outer training partition. Logistic calibration and all AlphaGenome/baseline stacks were trained on the selected inner out-of-fold EGrf probabilities and evaluated on the untouched outer cell. Selected hyperparameter frequencies and final metrics are in Supplementary Table 7.

#### Same-enhancer estimands

The conditional model uses frozen V1 deletion strength and rerun baseline activity. The conditional odds ratio estimates association after fixing an enhancer stratum and adjusting for the stated covariates. The predictive AUC analysis is different: it refits the conditional models with one enhancer held out and ranks only the candidate genes for that enhancer. Equal-enhancer AUC gives each enhancer weight 1; pair-weighted AUC weights by the number of positive-negative comparisons. Their divergence at high multiplicity is therefore expected and is reported explicitly.

The conditional odds ratio was 1.48 per standard deviation (enhancer-cluster sandwich 95% interval 0.97-2.23; robust Wald P=0.066). The one-degree-of-freedom likelihood-ratio P value was 0.034. These use different uncertainty assumptions; neither the smaller likelihood-ratio P value nor the borderline robust association establishes a predictive ranking gain.

#### Matching diagnostics

Standardized mean differences were calculated before and after matching for all five matching variables. Joint common support required each positive to lie within the negative range for all five variables. The sole caliper sensitivity used the 95th percentile of the original matching-distance distribution (1.74244); the threshold was not optimized against AlphaGenome outputs. Experimental class labels identify the two groups being matched.

#### Supporting K562 analyses

K562 inference first averaged pair differences within gene and then weighted genes equally. Robust estimands were calculated from the 121 gene-level contrasts. The top-tail removal analysis was descriptive and did not alter the primary cohort. Benchmark collision was evaluated by exact gene and interval joins on the recorded genome build and by same-gene interval overlap with ENCODE-rE2G. These results justify the supporting status of K562 and are not used as independent evidence for AstroREG.

### Supplementary Tables

#### Supplementary Table 1. AlphaGenome increment across the unified context design

| Context | Holdout | Context AP | + V1 AlphaGenome AP | AP increment (enhancer-cluster 95% interval) | Log-loss improvement (95% interval) |
|---|---|---|---|---|---|
| Linear | Enhancer | 0.3965 | 0.5340 | 0.1375 (0.0765-0.1964) | 0.01909 (0.00912-0.02982) |
| Linear | Gene + enhancer | 0.3648 | 0.5055 | 0.1406 (0.0760-0.1987) | 0.01785 (0.00771-0.02879) |
| Spline expression | Enhancer | 0.5875 | 0.6603 | 0.0728 (0.0306-0.1178) | 0.01658 (0.00704-0.02734) |
| Spline expression | Gene + enhancer | 0.5613 | 0.6310 | 0.0697 (0.0264-0.1144) | 0.01626 (0.00630-0.02754) |

In the linear enhancer-held-out specification, the standardized AlphaGenome coefficient was 0.9669 (enhancer-cluster 95% interval 0.5477-1.5565).

### Supplementary Table 2. Version-matched predicted-baseline control

| Context | Holdout | Context + rerun baseline AP | + rerun AlphaGenome AP | AP increment (enhancer-cluster 95% interval) | Log-loss improvement (95% interval) |
|---|---|---|---|---|---|
| Linear | Enhancer | 0.4014 | 0.5362 | 0.1347 (0.0734-0.1931) | 0.01849 (0.00841-0.02936) |
| Linear | Gene + enhancer | 0.3618 | 0.5101 | 0.1482 (0.0821-0.2028) | 0.01724 (0.00696-0.02824) |
| Spline expression | Enhancer | 0.5862 | 0.6579 | 0.0718 (0.0298-0.1164) | 0.01631 (0.00668-0.02706) |
| Spline expression | Gene + enhancer | 0.5592 | 0.6295 | 0.0703 (0.0274-0.1148) | 0.01581 (0.00578-0.02705) |

Rerun predicted baseline alone had raw AP 0.0694 and ROC AUC 0.4885. Figure 3b instead shows the fitted dual-holdout baseline-only model (out-of-fold AP 0.0534); the two values represent different prediction vectors. Rerun deletion strength and rerun baseline had Spearman rho 0.0050.

### Supplementary Table 3a. Distal sensitivity

| Population | Relations | Hits | Prevalence | Raw V1 AP (enhancer-cluster 95% interval) | Raw V1 ROC AUC (enhancer-cluster 95% interval) | Mean score difference (enhancer-cluster 95% interval) |
|---|---|---|---|---|---|---|
| Enhancer-target distance at least 10 kb | 2,243 | 101 | 0.0450 | 0.4195 (0.3223-0.5177) | 0.6821 (0.6026-0.7585) | 0.08095 (0.04808-0.12111) |

This post-hoc sensitivity uses the frozen score and outcome definitions and removes only relations with absolute enhancer-target distance below 10 kb. It was not used to redefine the primary cohort.

### Supplementary Table 3b. Request-window geometry sensitivities

| Population | Relations | Hits | Raw V1 AP | Raw V1 ROC AUC | Dual-holdout context AP | + V1 AlphaGenome AP | AP increment (95% interval) |
|---|---|---|---|---|---|---|---|
| Full cohort | 2,307 | 133 | 0.4785 | 0.7257 | 0.3648 | 0.5055 | 0.1406 (0.0760-0.1987) |
| Fully contained exon union | 2,271 | 129 | 0.4739 | 0.7318 | 0.3867 | 0.5150 | 0.1284 (0.0645-0.1900) |
| TSS at least 64 kb from boundary | 2,276 | 133 | 0.4788 | 0.7250 | 0.3987 | 0.5334 | 0.1347 (0.0739-0.1937) |

### Supplementary Table 4. Retrospective fixed-budget ranking

| Model | Precision@25 | Precision@50 | Precision@100 | Recall, top 1% | Recall, top 5% | Recall, top 10% |
|---|---|---|---|---|---|---|
| Dual-holdout context | 0.64 | 0.56 | 0.49 | 0.113 | 0.414 | 0.534 |
| Context + V1 AlphaGenome | 0.96 | 0.78 | 0.58 | 0.173 | 0.466 | 0.602 |
| Fully nested EGrf | 0.92 | 0.82 | 0.56 | 0.165 | 0.451 | 0.647 |
| Fully nested EGrf + V1 AlphaGenome | 0.92 | 0.86 | 0.69 | 0.165 | 0.534 | 0.662 |

Values summarize the completed cohort and must not be interpreted as prospective experimental yield.

### Supplementary Table 5. Same-enhancer ranking and candidate multiplicity

| Score | Multiplicity | Enhancers | Relations | Hits | Equal-enhancer AUC | Pair-weighted AUC |
|---|---|---|---|---|---|---|
| AlphaGenome, unadjusted | 2 candidates | 33 | 66 | 33 | 0.818 | 0.818 |
| AlphaGenome, unadjusted | 3-4 candidates | 44 | 154 | 50 | 0.780 | 0.813 |
| AlphaGenome, unadjusted | ≥5 candidates | 29 | 251 | 32 | 0.592 | 0.536 |
| Adjusted context, OOF | 2 candidates | 33 | 66 | 33 | 0.909 | 0.909 |
| Adjusted context, OOF | 3-4 candidates | 44 | 154 | 50 | 0.822 | 0.830 |
| Adjusted context, OOF | ≥5 candidates | 29 | 251 | 32 | 0.793 | 0.744 |
| Adjusted context + AlphaGenome, OOF | 2 candidates | 33 | 66 | 33 | 0.939 | 0.939 |
| Adjusted context + AlphaGenome, OOF | 3-4 candidates | 44 | 154 | 50 | 0.822 | 0.830 |
| Adjusted context + AlphaGenome, OOF | ≥5 candidates | 29 | 251 | 32 | 0.778 | 0.736 |

Overall adjusted context AUC was 0.8412 with equal enhancer weighting and 0.7823 with pair weighting. Adding AlphaGenome gave 0.8465 and 0.7797, respectively.

### Supplementary Table 6. Covariate balance in matching without AlphaGenome outputs

| Variable | Absolute SMD before | Absolute SMD after | Positive fraction in feature-specific support |
|---|---|---|---|
| Log distance | 1.280 | 0.144 | 0.992 |
| Log ABC score | 0.830 | 0.208 | 0.932 |
| Log enhancer length | 0.118 | 0.020 | 1.000 |
| Log measured expression | 0.149 | 0.150 | 0.880 |
| Log assayed-cell count | 0.057 | 0.006 | 1.000 |

The 133-pair mean difference was 0.07568 (95% interval 0.04854-0.10763). Joint common support covered 0.8195 of positives. The fixed caliper retained 126 pairs and gave 0.06654 (0.04064-0.09783). These historical 20,000-resample intervals are taken from the archived machine-readable matching_diagnostics_summary.json, included in source_data/; no matching analysis was rerun for this revision. Matching used the five context variables without AlphaGenome outputs. Residual absolute SMDs were 0.144 for distance, 0.208 for ABC and 0.150 for expression; matching therefore reduced some covariate differences but did not eliminate imbalance. Exact balance diagnostics are provided in source_data/matching_balance_diagnostics.csv.

### Supplementary Table 7. Fully nested EGrf model and tuning summary

| Model | AP | ROC AUC | Log loss | Brier score | Status |
|---|---|---|---|---|---|
| Released author EGrf probability | 0.5589 | 0.9002 | 0.1446 | 0.0366 | Descriptive; globally tuned before author cross-fitting |
| Fully nested EGrf, raw | 0.5372 | 0.8977 | 0.1409 | 0.0371 | Outer OOF |
| Fully nested EGrf, calibrated | 0.5475 | 0.8987 | 0.1410 | 0.0373 | Outer OOF |
| Fully nested EGrf + V1 AlphaGenome | 0.6137 | 0.9010 | 0.1306 | 0.0332 | Outer OOF |
| Fully nested EGrf + rerun baseline | 0.5411 | 0.8964 | 0.1422 | 0.0376 | Outer OOF |
| Fully nested EGrf + rerun baseline + rerun AlphaGenome | 0.6053 | 0.8970 | 0.1323 | 0.0336 | Outer OOF |

Selected hyperparameters across 25 outer cells were: `(mtry=2, extratrees, node=1)` in 6 cells; `(3, extratrees, 10)` in 6; `(2, extratrees, 10)` in 3; `(2, gini, 1)` in 3; `(3, gini, 1)` in 2; `(5, extratrees, 10)` in 2; and three other combinations in one cell each. Tuning was performed within each outer training partition to keep outer test cells excluded from hyperparameter selection.

### Supplementary Table 8. Robust K562 gene-equal estimands

| Population | Estimand | Estimate | Gene-bootstrap 95% interval | Genes |
|---|---|---|---|---|
| All | Mean | 0.4432 | 0.1355-0.7439 | 121 |
| All | 10% trimmed mean | 0.3351 | 0.1625-0.5627 | 121 |
| All | 20% trimmed mean | 0.1659 | 0.0715-0.3321 | 121 |
| All | 20% winsorized mean | 0.3252 | 0.1241-0.5435 | 121 |
| All | Median | 0.0235 | 0.0075-0.1202 | 121 |
| Gene-disjoint confirmation | Mean | 0.1012 | -0.4483-0.5278 | 24 |
| Gene-disjoint confirmation | Median | 0.0549 | 0.0038-0.3045 | 24 |

### Supplementary Table 9. K562 public-benchmark overlap

| Audit | Overlapping pairs | Total pairs | Fraction |
|---|---|---|---|
| Same Gasperini 2019 source study | 161 | 161 | 1.000 |
| Released AlphaGenome exact gene + hg38 enhancer | 111 | 161 | 0.689 |
| Released high-confidence metadata exact gene + hg19 candidate enhancer | 136 | 161 | 0.845 |
| ENCODE-rE2G same-gene hg38 interval overlap | 71 | 161 | 0.441 |
| ENCODE-rE2G overlap marked regulated | 68 | 161 | 0.422 |
| ENCODE-rE2G matched-negative overlap marked regulated | 5 | 161 | 0.031 |
| Strict regulated-positive/nonregulated-negative harmonized pair | 9 | 161 | 0.056 |

#### Supplementary Table 10. Alternate grouped-fold sensitivity

| Seed | Context | Holdout | Context AP | + V1 AlphaGenome AP | AP increment | Log-loss improvement |
|---|---|---|---|---|---|---|
| 20260829 | Linear | Enhancer | 0.3939 | 0.5281 | 0.1342 | 0.01616 |
| 20260829 | Linear | Gene + enhancer | 0.3928 | 0.5260 | 0.1331 | 0.01670 |
| 20260829 | Spline expression | Enhancer | 0.5884 | 0.6493 | 0.0610 | 0.01299 |
| 20260829 | Spline expression | Gene + enhancer | 0.5738 | 0.6315 | 0.0577 | 0.01154 |
| 20260830 | Linear | Enhancer | 0.4014 | 0.5339 | 0.1325 | 0.01780 |
| 20260830 | Linear | Gene + enhancer | 0.3982 | 0.5311 | 0.1329 | 0.01720 |
| 20260830 | Spline expression | Enhancer | 0.5993 | 0.6596 | 0.0603 | 0.01425 |
| 20260830 | Spline expression | Gene + enhancer | 0.5882 | 0.6517 | 0.0635 | 0.01404 |
| 20260831 | Linear | Enhancer | 0.3916 | 0.5312 | 0.1396 | 0.01724 |
| 20260831 | Linear | Gene + enhancer | 0.3727 | 0.5139 | 0.1412 | 0.01698 |
| 20260831 | Spline expression | Enhancer | 0.5867 | 0.6535 | 0.0668 | 0.01380 |
| 20260831 | Spline expression | Gene + enhancer | 0.5686 | 0.6397 | 0.0712 | 0.01397 |

### Atlas extension: exploratory supplementary methods and results

Tables 1-10 describe the deletion analysis; Tables 11-19 describe the exploratory Atlas extension and its controls. The analyses have different estimands and resampling counts.

#### Analysis sequence and interpretation

The study used an assay-ascertained cohort whose outcomes were already known during the Atlas extension. Local analysis freezes record the order of decisions within that exploratory work; they are not external preregistration or an independent confirmatory cohort.

| Analysis | Recorded order of specification | Interpretation |
|---|---|---|
| Nine-SNV Atlas summaries and initial standalone contrasts | Mean absolute RNA response, secondary signed response and AVI were specified before cohort metrics; initial contrasts compared Atlas magnitude with signed V1 and rerun deletion | Exploratory, with previous exposure to AstroREG outcomes |
| Absolute-deletion standalone controls | Added after the initial Atlas analysis to match direction handling | Post-hoc magnitude comparison; original signed results retained |
| Equal-weight Atlas fusion | Fixed after standalone Atlas results and before this fusion module was computed | No weights fitted to outcomes, but the choice of rule followed earlier results |
| Absolute-deletion fusion control | Fixed after the Atlas fusion results and before the control was computed | Post-result comparison of alternative fusion partners |
| Full deletion, shuffle and middle-third encodings | Defined before full-cohort queries, following a three-locus technical pilot | Exploratory comparison of computational perturbations |
| Magnitude summaries of those encodings | Added during full-cohort inference after the Atlas magnitude results, before inspecting these encoding metrics | Outcome-informed secondary analysis; all encodings retained |
| AVI-augmented comparators | Fixed before AVI-augmented fits and complete-cohort handoff, using the existing folds and seven-input model specifications | Exploratory enhancer-level feature augmentation |

These stages are documented in the archived configurations atlas_pilot.json, atlas_statistics.json, atlas_magnitude_controls.json, preassay_atlas_fixed_plan.json, preassay_atlas_magnitude_fixed_plan.json, perturbation_cohort.json and preassay_avi_plan.json. Their original counts and paired resampling schemes are retained below.

### Supplementary Table 11. All fixed Atlas fusion rules

| Comparator | Rule | AP | ROC AUC |
|---|---|---|---|
| Logistic | Absolute deletion fusion | 0.5636 | 0.9102 |
| Logistic | Atlas magnitude fusion | 0.6155 | 0.9179 |
| Logistic | Atlas signed-decrease fusion | 0.3323 | 0.7608 |
| Logistic | Baseline rank | 0.5110 | 0.9070 |
| Random forest | Absolute deletion fusion | 0.5720 | 0.9014 |
| Random forest | Atlas magnitude fusion | 0.6357 | 0.9094 |
| Random forest | Atlas signed-decrease fusion | 0.3410 | 0.7406 |
| Random forest | Baseline rank | 0.5157 | 0.8915 |

All 2,307 relations and 133 decreasing hits. Base and response scores use training-reference mid-CDF ranks; fusion weights are equal. Scores are not probabilities. Magnitude and adverse signed rules are both shown. Marginal intervals and baseline comparisons are supplied in source_data/fusion_marginal_intervals.csv and atlas_baseline_contrasts.csv.

### Supplementary Table 12. Direct Atlas-minus-absolute-deletion fusion comparisons

| Comparator | Metric | Resampling unit | Increment | 95% CI |
|---|---|---|---|---|
| Logistic | AP | Enhancer | 0.0519 | [0.0098, 0.0957] |
| Logistic | ROC AUC | Enhancer | 0.0077 | [-0.0111, 0.0272] |
| Random forest | AP | Enhancer | 0.0637 | [0.0220, 0.1094] |
| Random forest | ROC AUC | Enhancer | 0.0079 | [-0.0116, 0.0286] |
| Logistic | AP | Gene | 0.0519 | [0.0065, 0.0998] |
| Logistic | ROC AUC | Gene | 0.0077 | [-0.0109, 0.0285] |
| Random forest | AP | Gene | 0.0637 | [0.0143, 0.1136] |
| Random forest | ROC AUC | Gene | 0.0079 | [-0.0111, 0.0288] |

Each interval uses 5,000 paired resamples of frozen predictions. Enhancer and gene clustering are alternative dependence sensitivities, not a joint two-way bootstrap or independent replication. The wider outcome-informed research history and model-refitting variability are not included.

### Supplementary Table 13. Snapshot-matched association and fixed ranking

| Deletion snapshot | Odds ratio per SD | Robust 95% CI | Robust Wald P |
|---|---|---|---|
| V1 | 1.476 | [0.975, 2.235] | 0.0660 |
| Matched rerun | 1.476 | [0.971, 2.244] | 0.0684 |

Models use 471 relations, 106 mixed-outcome enhancers and 115 decreasing hits; the five context covariates and Efron likelihood match the historical specification. Reported intervals and P values are enhancer-cluster sandwich estimates, distinct from model-based likelihood tests. The fixed-score ranking table is source_data/matched_fixed_ranking.csv; its historical 20,000 resamples are not the 5,000 used for Atlas fusion. The matched learned LOEO prediction was not performed and is not inferred from these association or fixed-ranking results.

### Supplementary Table 14. Within-enhancer scores and paired comparisons

| Score or paired contrast | Equal-enhancer AUC or increment [95% CI] | Pair-weighted AUC or increment [95% CI] |
|---|---|---|
| Signed deletion (rerun) | 0.7556 [0.6779, 0.8303] | 0.6658 [0.5525, 0.7792] |
| Signed deletion (V1) | 0.7406 [0.6620, 0.8160] | 0.6380 [0.5324, 0.7474] |
| Atlas signed decrease | 0.4708 [0.3830, 0.5605] | 0.4582 [0.3401, 0.5849] |
| Atlas magnitude | 0.8965 [0.8443, 0.9422] | 0.8987 [0.8373, 0.9462] |
| Absolute deletion (V1) | 0.8101 [0.7432, 0.8733] | 0.7924 [0.7057, 0.8709] |
| Absolute deletion (rerun) | 0.8861 [0.8358, 0.9318] | 0.8076 [0.7255, 0.8905] |
| Proximity | 0.8382 [0.7777, 0.8941] | 0.7772 [0.7083, 0.8504] |
| ABC | 0.8213 [0.7600, 0.8778] | 0.7532 [0.6843, 0.8310] |
| Atlas magnitude minus absolute deletion (V1) | 0.0864 [0.0158, 0.1571] | 0.1063 [0.0138, 0.1958] |
| Atlas magnitude minus absolute deletion (rerun) | 0.0105 [-0.0451, 0.0648] | 0.0911 [-0.0089, 0.1831] |
| Atlas magnitude minus proximity | 0.0584 [-0.0094, 0.1250] | 0.1215 [0.0230, 0.2086] |

All rows use the same 106 mixed-outcome enhancers, 471 relations, 115 decreasing hits and 395 hit-comparison pairs; intervals use 20,000 enhancer resamples, paired for differences. Equal-enhancer AUC weights each enhancer equally, whereas pair-weighted AUC gives more weight to enhancers with more hit-comparison pairs. The Atlas-minus-proximity interval crosses zero under equal-enhancer weighting but excludes zero under pair weighting; neither is evidence of equivalence. Strong standalone ranking does not by itself establish an increment over proximity or a fitted context model. Magnitude scores and contrasts are from source_data/within_enhancer.csv, signed scores from atlas_signed_within_enhancer.csv and ABC from perturbation_metrics.csv; full-span sensitivities remain in within_enhancer.csv. Separate context-plus-one-exposure conditional fits are in source_data/magnitude_conditional.csv. Atlas OR=4.971, enhancer-robust 95% interval [1.515, 16.312], nominal P=0.0082. These post-hoc fits do not test Atlas beyond an included absolute-deletion score. Absolute deletion and shuffle ORs were 1.506 and 1.398, with robust intervals crossing one. Differences in P values are not tests of exposure superiority.

### Supplementary Table 15. External WTC11 MHC fixed-score stress test

| Score or contrast | Correlation or increment | 95% CI | CREs | Relations |
|---|---|---|---|---|
| Atlas signed decrease | -0.0067 | [-0.0438, 0.0303] | 317 | 5557 |
| Atlas magnitude | 0.1162 | [0.0841, 0.1479] | 317 | 5557 |
| rE2G score | 0.1142 | [0.0811, 0.1480] | 317 | 5557 |
| Signed Atlas fusion | 0.0661 | [0.0318, 0.0998] | 317 | 5557 |
| Atlas magnitude fusion | 0.1297 | [0.0970, 0.1629] | 317 | 5557 |
| Signed fusion minus rE2G | -0.0480 | [-0.0761, -0.0200] | 317 | 5557 |
| Magnitude fusion minus rE2G | 0.0155 | [-0.0074, 0.0372] | 317 | 5557 |

rE2G score denotes the ENCODE-rE2G continuous relation score (the source CSV column named Score). Endpoint is negative median published guide log2 fold change; estimates are equal-CRE mean within-CRE Spearman correlations or direct increments. Intervals use 10,000 CRE-summary resamples within one locus and donor, not independent biological replication. All complete-score, full-v46-span, FRACTEL and equal-gene analyses are in source_data/mhc_all_results.csv. The equal-gene incremental comparator is within-CRE-rank rE2G, not raw rE2G. One incomplete relation remains missing. The primary 317-CRE denominator excludes 11 complete-score CREs with fewer than four candidates, not CREs selected for accuracy.

### Supplementary Table 16. Retained secondary controls and execution identity

| Control family | Frozen complete table | Interpretation |
|---|---|---|
| Scalar AVI-augmented pre-assay models | avi_metrics.csv; avi_contrasts.csv | Enhancer-level AVI cannot order genes within one enhancer; not RNA-output predictive fitting |
| Historical/new comparator probabilities and rank mapping | preassay_all_metrics.csv | New sklearn comparators differ from historical ranger; normalization is not an Atlas increment |
| Full deletion, dinucleotide shuffle, middle-third deletion | perturbation_metrics.csv; perturbation_contrasts.csv | Encoding differences retained; not equivalent biological interventions |
| Snapshot equality | perturbation_cohort_analysis_summary.json | All 2,307 fresh full-deletion scores equal cached rerun; server weight identity unknown |

All filenames in this section refer to atlas_extension/source_data/ in the public reproducibility package. Complete rule families, including uncertain or adverse results, are retained. The operation-specific version/use record is provided in docs/ATLAS_VERSION_AND_PROVENANCE.md in the public reproducibility package (version 1.0.0; https://doi.org/10.5281/zenodo.23351607). No new RNA-output-trained predictive model or matched LOEO fit was executed during the extension; historical approved fitted analyses retain their own identity. Exact machine-precision values and identifiers remain unchanged in the accompanying CSV tables.

### Supplementary Table 17. Standalone relation-level scores and direct paired contrasts

| Score | AP [95% CI] | ROC AUC [95% CI] | Source |
|---|---|---|---|
| Proximity | 0.3075 [0.2352, 0.3973] | 0.8105 [0.7679, 0.8510] | A |
| ABC | 0.4013 [0.3162, 0.4908] | 0.7873 [0.7374, 0.8358] | C |
| Signed deletion (V1) | 0.4785 [0.3915, 0.5659] | 0.7257 [0.6615, 0.7882] | A |
| Signed deletion (rerun) | 0.4848 [0.3985, 0.5718] | 0.7405 [0.6775, 0.8018] | A |
| Absolute deletion (V1) | 0.4915 [0.4032, 0.5794] | 0.8324 [0.7871, 0.8751] | B |
| Absolute deletion (rerun) | 0.5018 [0.4140, 0.5892] | 0.8581 [0.8187, 0.8941] | B |
| Atlas signed decrease | 0.2666 [0.1900, 0.3500] | 0.5232 [0.4486, 0.5951] | A |
| Atlas magnitude | 0.5570 [0.4713, 0.6413] | 0.8712 [0.8310, 0.9075] | A |
| AVI (enhancer-level) | 0.0683 [0.0554, 0.0867] | 0.5504 [0.5004, 0.6004] | A |

| Paired contrast | AP increment [95% CI] | ROC-AUC increment [95% CI] | Source |
|---|---|---|---|
| Atlas magnitude minus signed deletion (V1) | 0.0785 [0.0101, 0.1495] | 0.1455 [0.0801, 0.2125] | A |
| Atlas magnitude minus signed deletion (rerun) | 0.0722 [0.0041, 0.1429] | 0.1306 [0.0661, 0.1970] | A |
| Atlas signed decrease minus signed deletion (V1) | -0.2119 [-0.3030, -0.1177] | -0.2025 [-0.2979, -0.1097] | A |
| Atlas signed decrease minus signed deletion (rerun) | -0.2182 [-0.3082, -0.1246] | -0.2173 [-0.3136, -0.1238] | A |
| Atlas magnitude minus absolute deletion (V1) | 0.0656 [0.0006, 0.1341] | 0.0388 [-0.0031, 0.0821] | B |
| Atlas magnitude minus absolute deletion (rerun) | 0.0552 [-0.0098, 0.1225] | 0.0130 [-0.0274, 0.0542] | B |
| Atlas magnitude minus proximity | 0.2495 [0.1573, 0.3306] | 0.0607 [0.0149, 0.1064] | B |

All rows use 2,307 relations, 745 enhancers and 133 decreasing hits (prevalence 0.0577). Proximity ranks shorter absolute enhancer-target distances higher. Signed summaries rank predicted decreases higher; absolute summaries discard sign. AVI is constant across candidate genes of an enhancer. These are standalone scores, without fusion or supervised score calibration.

Intervals are nominal 95% percentile intervals from 20,000 enhancer resamples; paired contrasts use the same weights for both scores. Source A is source_data/atlas_standalone_metrics.csv (the original atlas_fixed_score_metrics.csv); B is atlas_magnitude_control_metrics.csv, restricted to complete_9_SNV and enhancer_id; C is perturbation_metrics.csv. Marginal intervals retain each module's recorded draws, so the same point estimate can have slightly different interval endpoints in another module. The Atlas-magnitude marginal interval from B is [0.4725, 0.6396] for AP and [0.8317, 0.9071] for ROC AUC. Direct paired intervals, rather than subtraction of marginal limits, support comparisons. Initial Atlas contrasts and later absolute-deletion controls have the different analysis statuses stated above. The rerun magnitude-matched AP and ROC-AUC intervals cross zero; the V1 AP interval does not. The snapshot-specific comparisons are therefore reported separately; comparisons with supervised-comparator point estimates are descriptive.

### Supplementary Table 18. Perturbation-encoding controls

| Perturbation and summary | AP [95% CI] | ROC AUC [95% CI] | Equal-enhancer AUC [95% CI] |
|---|---|---|---|
| Full deletion, signed | 0.4848 [0.3985, 0.5718] | 0.7405 [0.6775, 0.8018] | 0.7556 [0.6779, 0.8303] |
| Dinucleotide shuffle, signed | 0.5104 [0.4244, 0.5971] | 0.7849 [0.7265, 0.8407] | 0.8211 [0.7512, 0.8846] |
| Middle-third deletion, signed | 0.4126 [0.3238, 0.5046] | 0.6648 [0.5892, 0.7386] | 0.6839 [0.6000, 0.7637] |
| Full deletion, magnitude | 0.5018 [0.4128, 0.5911] | 0.8581 [0.8179, 0.8946] | 0.8861 [0.8358, 0.9318] |
| Dinucleotide shuffle, magnitude | 0.5234 [0.4346, 0.6116] | 0.8707 [0.8322, 0.9058] | 0.8920 [0.8410, 0.9379] |
| Middle-third deletion, magnitude | 0.4734 [0.3816, 0.5695] | 0.8484 [0.8061, 0.8878] | 0.8486 [0.7890, 0.9035] |

| Paired contrast | AP increment [95% CI] | ROC-AUC increment [95% CI] | Equal-enhancer AUC increment [95% CI] |
|---|---|---|---|
| Shuffle minus full deletion, signed | 0.0256 [-0.0067, 0.0559] | 0.0444 [-0.0010, 0.0907] | 0.0655 [0.0095, 0.1235] |
| Middle-third minus full deletion, signed | -0.0721 [-0.1243, -0.0199] | -0.0757 [-0.1386, -0.0150] | -0.0717 [-0.1503, 0.0053] |
| Shuffle minus full deletion, magnitude | 0.0215 [-0.0058, 0.0493] | 0.0126 [-0.0187, 0.0456] | 0.0059 [-0.0370, 0.0493] |
| Middle-third minus full deletion, magnitude | -0.0284 [-0.0777, 0.0248] | -0.0097 [-0.0456, 0.0270] | -0.0374 [-0.0963, 0.0180] |

Cohort-level AP and ROC AUC use all 2,307 relations and 133 hits; within-enhancer AUC uses the 106 mixed-outcome enhancers, 471 relations and 115 hits. Intervals use 20,000 paired enhancer resamples for differences. Full deletion and cached client-0.7.0 rerun scores were identical across all 2,307 relations; their zero contrasts remain in source_data/perturbation_contrasts.csv. Point estimates, marginal intervals and pair-weighted sensitivities are in perturbation_metrics.csv; all direct differences are in perturbation_contrasts.csv.

The deterministic dinucleotide shuffle preserves sequence length, whereas both deletion encodings shorten the sequence. The signed shuffle-minus-deletion AP interval crosses zero, while its equal-enhancer AUC interval excludes zero. With magnitude matching, the corresponding AP and equal-enhancer intervals both cross zero. Thus useful response ranking also occurs under length-preserving sequence disruption, but these encodings do not isolate coordinate displacement from all other perturbation differences or establish a biological mechanism. The signed comparisons were specified before full-cohort queries; the magnitude comparisons were added after Atlas results and remain secondary exploratory analyses.

### Supplementary Table 19. Enhancer-level AVI augmentation

| Comparator | Base AP | AVI-augmented AP | Base ROC AUC | AVI-augmented ROC AUC |
|---|---|---|---|---|
| Logistic | 0.5106 | 0.5135 | 0.9066 | 0.9051 |
| Random forest | 0.5153 | 0.5185 | 0.8908 | 0.8949 |

| Comparator | Resampling unit | AP increment [95% CI] | ROC-AUC increment [95% CI] | Log-loss improvement [95% CI] |
|---|---|---|---|---|
| Logistic | Enhancer | 0.0029 [-0.0050, 0.0113] | -0.0015 [-0.0040, 0.0007] | -0.0001 [-0.0010, 0.0008] |
| Logistic | Gene | 0.0029 [-0.0055, 0.0131] | -0.0015 [-0.0039, 0.0010] | -0.0001 [-0.0011, 0.0008] |
| Random forest | Enhancer | 0.0031 [-0.0148, 0.0204] | 0.0042 [-0.0035, 0.0139] | 0.0106 [-0.0019, 0.0277] |
| Random forest | Gene | 0.0031 [-0.0149, 0.0210] | 0.0042 [-0.0044, 0.0143] | 0.0106 [-0.0018, 0.0283] |

All 2,307 relations, 745 enhancers, 1,054 genes and 133 hits use the same 25 crossed outer cells as the seven-input comparators. A single mean raw AVI score over the nine fixed SNVs was appended before nested fitting, with the existing model classes, grids, folds and transforms. These are out-of-fold model probabilities, distinct from the training-reference ranks used in Table 11, whose AP values can differ slightly because the fold-specific rank mapping changes pooled ordering. Intervals use 5,000 paired enhancer or gene resamples of fixed out-of-fold predictions; AP and ROC-AUC increments are augmented minus base, whereas log-loss improvement is base minus augmented. Source files are source_data/avi_metrics.csv and avi_contrasts.csv.

No AVI-augmentation interval shown excludes zero. This result does not demonstrate that the RNA-response increment must arise from gene-specific information: AVI is a different enhancer-level summary, and absence of an established increment is not a causal mediation test. By construction, standalone AVI ties all genes within an enhancer and has within-enhancer AUC 0.5. The AVI analysis was locally frozen before augmented fits but used a cohort whose outcomes were already known.

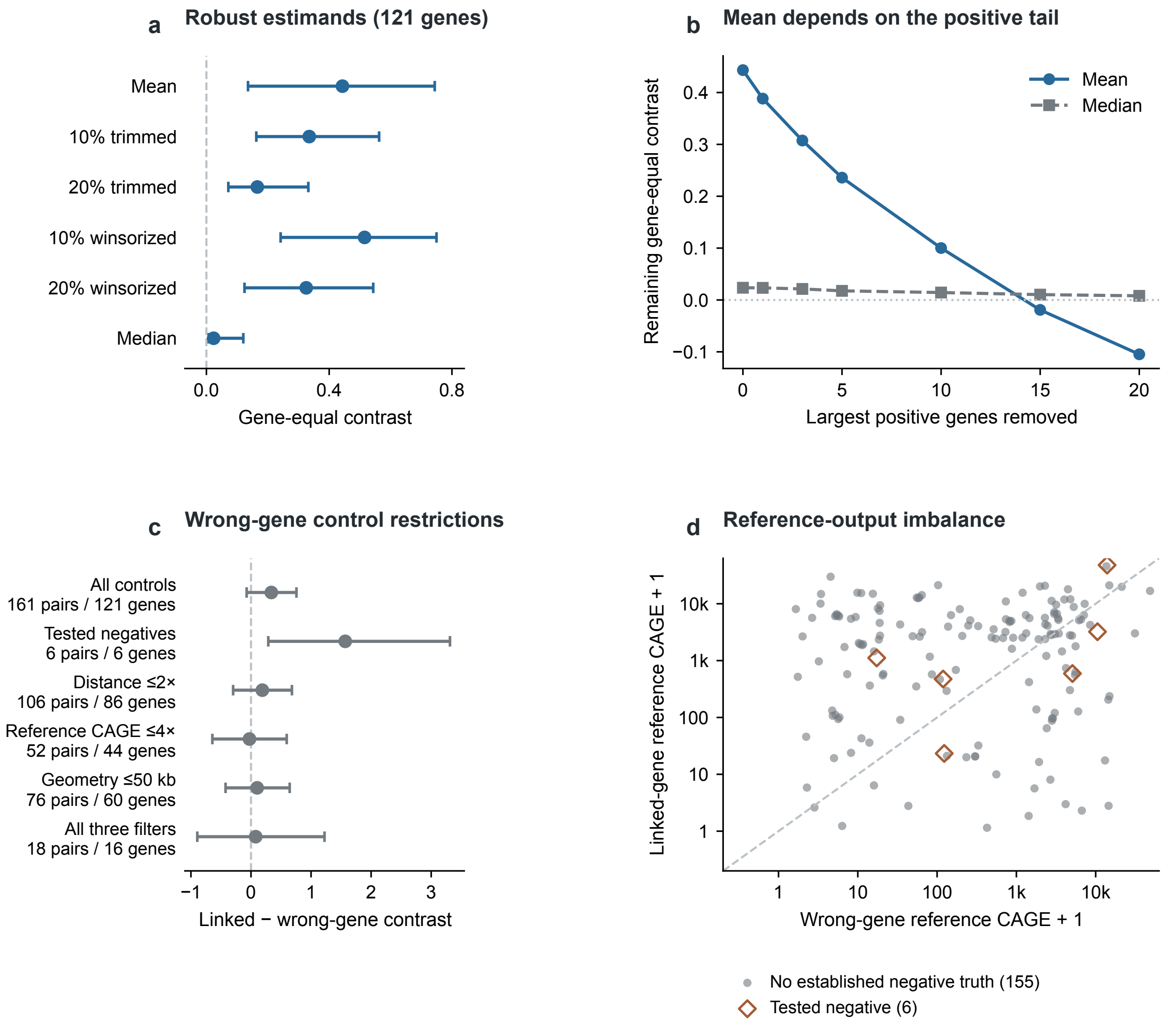


**Supplementary Figure 1 | Supporting K562 robustness and wrong-gene control diagnostics. a,** Gene-equal positive-minus-matched-negative deletion-strength estimands under all six mean, trimmed, winsorized and median summaries (161 pairs, 121 genes); bars are 95% gene-bootstrap intervals from 20,000 resamples. **b,** Descriptive removal of the largest positive gene contrasts: blue circles/solid line, mean; gray squares/dashed line, median. Each point uses 121 minus the indicated number of removed genes, and no interval is plotted. **c,** Gene-equal linked-minus-wrong-gene contrasts for six original control groups, including all controls; points are means and bars are 95% gene-bootstrap intervals from the original 20,000 resamples. Pairs and genes are printed beside each group. **d,** Reference CAGE output for linked and wrong genes in all 161 comparisons after adding one on logarithmic axes (k = 1,000): six explicitly tested negatives are open brown diamonds, and 155 controls without established negative experimental truth are gray circles. The diagonal is equality. Baseline and request-geometry differences limit target-specific interpretation. All-cohort and gene-disjoint confirmation summaries remain in Supplementary Table 8; development and complete robust summaries remain in machine-readable K562 results. Benchmark overlap remains in Supplementary Table 9. K562 is supporting evidence, not independent validation.
